# No enhancement of soil carbon persistence by sheep grazing in a long-term calcareous grassland experiment

**DOI:** 10.1101/2024.09.11.611980

**Authors:** David Encarnation, Deborah Ashworth, Richard Bardgett, Mona Edwards, Clive Hambler, Jeppe Kristensen, Andy Hector

## Abstract

Soils hold a globally important carbon pool that is generally more persistent than the carbon stored in plant biomass. However, this carbon is becoming increasingly vulnerable to disturbances such as soil warming, fire, and erosion. Managing land to increase soil carbon sequestration and persistence may therefore improve long-term soil carbon storage and contribute to climate change mitigation. It has been hypothesized that grazing by large herbivores may enhance the persistence of soil carbon by increasing the amount of soil organic matter forming more stable associations with mineral particles (mineral-associated organic matter). We compared sheep-grazed and ungrazed plots within the Gibson Grazing and Successional Experiment located in the Upper Seeds calcareous grassland in Wytham Woods, Oxfordshire, using organic matter fractionation to estimate the surface (0-5 cm) carbon stocks in the mineral-associated and particulate organic matter fractions. Counter to predictions, after 35 years sheep grazing had not increased mineral-associated organic matter carbon stocks relative to ungrazed plots. We hypothesize that this indicates the saturation of mineral surfaces in both grazed and ungrazed treatments and the inability of grazing to increase soil nitrogen stocks and decrease pH to levels conducive for mineral-associated carbon sequestration. Only one of twelve soil properties examined showed statistically detectable responses to grazing: spring-grazing increased the C:N ratio in the mineral-associated organic matter. While the number of tests performed (24) means this may be a false-positive result, if genuine it would be consistent with a more direct pathway from plant exudates to mineral-associated organic matter formation due to compensatory growth in response to spring-grazing. Overall, the results of this long-term experiment do not support the hypothesis that grazing can improve the persistence of the soil carbon pool.

## Introduction

The global soil carbon pool is more than three times the size of the atmospheric carbon pool. Consequently, soil carbon sequestration (“transferring atmospheric CO_2_ into long-lived pools and storing it securely so it is not immediately reemitted”) is a useful tool to combat climate change (Lal, 2004). Land management regimes are often suggested as integral not only to soil carbon storage, but also to the long-term persistence of the soil carbon pool.

The amount of carbon stored in soils reflects the balance of inputs and outputs of carbon. When decomposition is not severely constrained by environmental conditions (mainly pH or oxygen availability), carbon storage is the combined result of the productivity of the system, the decomposability of the carbon inputs (intrinsic organic matter characteristics), and the protection against decomposition by physico-chemical interactions with mineral particles. Traditionally, intrinsic organic matter characteristics, such as lignin concentration, were considered the main drivers of decomposability. However, an emerging view suggests that the main driver of decomposability is “bioaccessibility”, i.e., how accessible the organic matter is to microbial decomposition (Just *et al*., 2021; Lehmann & Kleber, 2015). While the intrinsic characteristics of organic matter are still relevant (particularly for determining the initial decomposability), this bioaccessibility is largely driven by physical stabilization within soil aggregates and microaggregates as well as chemical stabilization via association with mineral particles (Six *et al*., 2002). This view divides soil organic matter into different fractions with different formation pathways, capacities for carbon storage, and levels of persistence in the soil (Cotrufo *et al*., 2019; Lavallee *et al.,* 2019; Von Lutzow *et al*., 2007).

One main pool of soil organic matter is **particulate organic matter (POM)**, which refers to the fraction of soil organic matter mostly composed of undecomposed plant structural compounds such as root and shoot structural tissues. Particulate organic matter is predominantly stabilized through the intrinsic characteristics of organic matter mentioned above, which can be less susceptible to microbial decomposition (Cotrufo *et al*., 2019). However, particulate organic matter is vulnerable to increases in decomposition driven by environmental changes such as soil warming (Abramoff *et al*., 2021). Thus, these intrinsic characteristics can only protect the particulate organic matter from decomposition for a few years to decades, though some types of particulate organic matter (e.g., when occluded in soil aggregates) can persist for hundreds of years (Angst *et al.,* 2023; Lugato *et al*., 2021; Wasak & Drewnik, 2015).

A second main pool is **mineral-associated organic matter (MAOM)**. This fraction of soil organic matter consists primarily of belowground carbon inputs such as root exudates. Whereas the stability of particulate organic matter relies on its intrinsic characteristics, mineral-associated organic matter derives its low bioavailability from this combination of physical protection in soil aggregates and microaggregates as well as chemical protection via sorption to mineral surfaces (Kristensen *et al*., 2022). This “physicochemical inhibition” renders mineral-associated organic matter more resistant to microbial decomposition and less vulnerable to physical and environmental perturbations compared to particulate organic matter (Lugato *et al*., 2021; Wasak & Drewnik, 2015). Mineral-associated organic matter can persist in soils for centuries, making it the main driver of long-term carbon storage in soils (Kristensen *et al*., 2022). As such, carbon sequestration efforts should consider not only carbon pool sizes but also persistence when prioritizing land use and management strategies.

Soil carbon stocks are highly sensitive to land and soil management, particularly grazing (Conant *et al*., 2017; Jones & Donnelly, 2004). Grasslands cover 20-30% of the Earth’s ice-free land area (Gibson & Newman, 2019; Wilson *et al*., 2018), and 36% of the UK’s land cover (Ward *et al*., 2016). They are important for the diversity of ecosystem services they provide, including food and forage production, tourism, recreation, biofuel production, and carbon storage (Gibson & Newman, 2019). Temperate grasslands such as those in the UK store over 12% of global carbon and are the third largest store of carbon in soil and vegetation (Ward *et al*., 2016). However, some grassland types are increasingly threatened globally due to a combination of land use change, agricultural intensification, climate change, and management changes such as fire suppression and land abandonment (Gibson & Newman, 2019). Global estimates suggest that 49% of grassland area has already been degraded often with associated declines in soil carbon stocks (Bardgett *et al*, 2021). As environmental and anthropogenic perturbations of grasslands become more common, we must understand the effects of grassland management practices on the resilience of soil carbon to perturbations.

Kristensen *et al*. (2022) hypothesized that grazing by large herbivores could enhance soil carbon persistence in grassy ecosystems by increasing the carbon stored in the more stable mineral-associated organic matter fraction. They suggest that large herbivores may achieve this through: (1) “increasing ecosystem metabolism and fertility”, (2) “favoring herbaceous vegetation with high belowground inputs and dense root nets”, and/or (3) “soil mixing and processing by large animals and associated fauna”. First, increasing fertility tends to lead to an increase in plant productivity. This increases the input of organic matter to the soil and enhances microbial biomass production, which leads to an increase in mineral-associated organic matter formation. Second, grazing can enhance the presence of fast-growing plant species that invest more resources in root biomass in deeper soil horizons and often have high rates of root exudation, which is key to the formation of mineral-associated organic matter. This creates more opportunities for these carbon inputs to form stable microaggregates and associate with mineral surfaces. Lastly, some large herbivores and their associated fauna mix the soil vertically, putting the organic matter inputs from the soil surface in close contact with the deeper mineral soil, which can lead to greater mineral-associated organic matter formation.

The formation of particulate organic matter and mineral-associated organic matter depends on many soil properties. First, all soil organic matter contains nitrogen and is produced via microbial processes that require nitrogen. As a result, soil carbon stocks depend on nitrogen availability (Cotrufo *et al*., 2019). Particulate organic matter tends to have a low nitrogen content (i.e., a high carbon to nitrogen ratio), and mineral-associated organic matter a relatively higher nitrogen content (i.e., a lower carbon to nitrogen ratio) (Cotrufo *et al*., 2019; Lugato *et al*., 2021). So, while mineral-associated organic matter is more stable long-term than particulate organic matter, its formation requires more nitrogen (Lugato *et al*., 2021). Second, pH influences carbon storage in both particulate and mineral-associated organic matter pools. Both particulate and mineral-associated organic matter carbon sequestration increases as the soil pH decreases. In the case of particulate organic matter, this is the result of reduced microbial decomposition at a lower pH (Lugato *et al*., 2021). For mineral-associated organic matter, sorption (the association of organic matter with mineral surfaces) increases as the pH decreases, with Lutzow *et al*. (2006) estimating that the maximum sorption occurs between a pH of 4.3-4.7. Within this pH range, ligand exchange between organic matter and compounds such as iron-, aluminum-, or manganese-oxides is maximized. At higher pH, there is competition for these exchanges from anions like CO_3_^2-^ and PO_3_^3-,^ whereas at lower pH values the oxides themselves start dissolving and become unavailable. Organic matter can become sequestered in the interlayers of expandable clays only at pH <5 (Lutzow *et al*., 2006). Finally, soil depth is an important factor, with the input of organic matter decreasing with depth and the carbon stored in deeper layers disproportionately stored in the mineral-associated fraction (given the increased abundance of mineral surfaces in deeper layers).

There is currently a lack of data from long-term designed experiments examining the effects of large herbivores – including domesticated species - on the distribution of grassland soil carbon into the different fractions of organic matter. This study used a long-term sheep grazing experiment on a calcareous grassland to test the primary hypothesis that grazing by large herbivores will enhance surface (0-5 cm) soil carbon persistence by increasing the amount of mineral-associated organic carbon. We also tested the associated hypothesis that grazing by large herbivores should increase soil fertility by determining the effects of sheep grazing on soil nitrogen stocks and C:N ratios, which would be expected to increase (stocks) and decrease (ratios) respectively over time.

## Materials and Methods

### Study site

Soil samples were collected from the long-term Gibson Grazing and Successional Experiment (‘Gibson experiment’, formerly known as the ‘Upper Seeds Experiment’) within the Upper Seeds grassland in the Wytham Woods estate (51°46’10.6” N, 1°19’54.9” W, 160 m above mean sea level) (Gibscon, CWD. 2011). The site lies on top of a small hill with a south-easterly aspect. Upper Seeds is a calcareous grassland approximately 10 ha in size (Figure 1) with a shallow (10-20 cm) soil crust on top of a calcareous bedrock made up of Jurassic coral reef (Stone, 2020). The site was in arable production from at least the Second World War through ca. 1980, when it was taken out of production and set aside for conservation purposes (Gibson, 1986). The resulting Upper Seeds soil is a shallow, well-drained calcareous clay soil recovering from the previous disturbance (including ploughing for arable production) (Taylor *et al*., 2011). The Gibson Grazing and Successional Experiment ran for 35 years (1985-2020). The experiment ran consistently until 2014 and had a few unsystematic gaps in grazing treatments until the end of the experiment in 2020. In 2021 fences between plots were removed so that sheep were free to graze the whole experiment for one growing season after the formal end of the experiment (see discussion). Soil samples were collected in January 2022 to assess the cumulative effects of 35 years of differences in grazing treatments.

**Figure 1.**
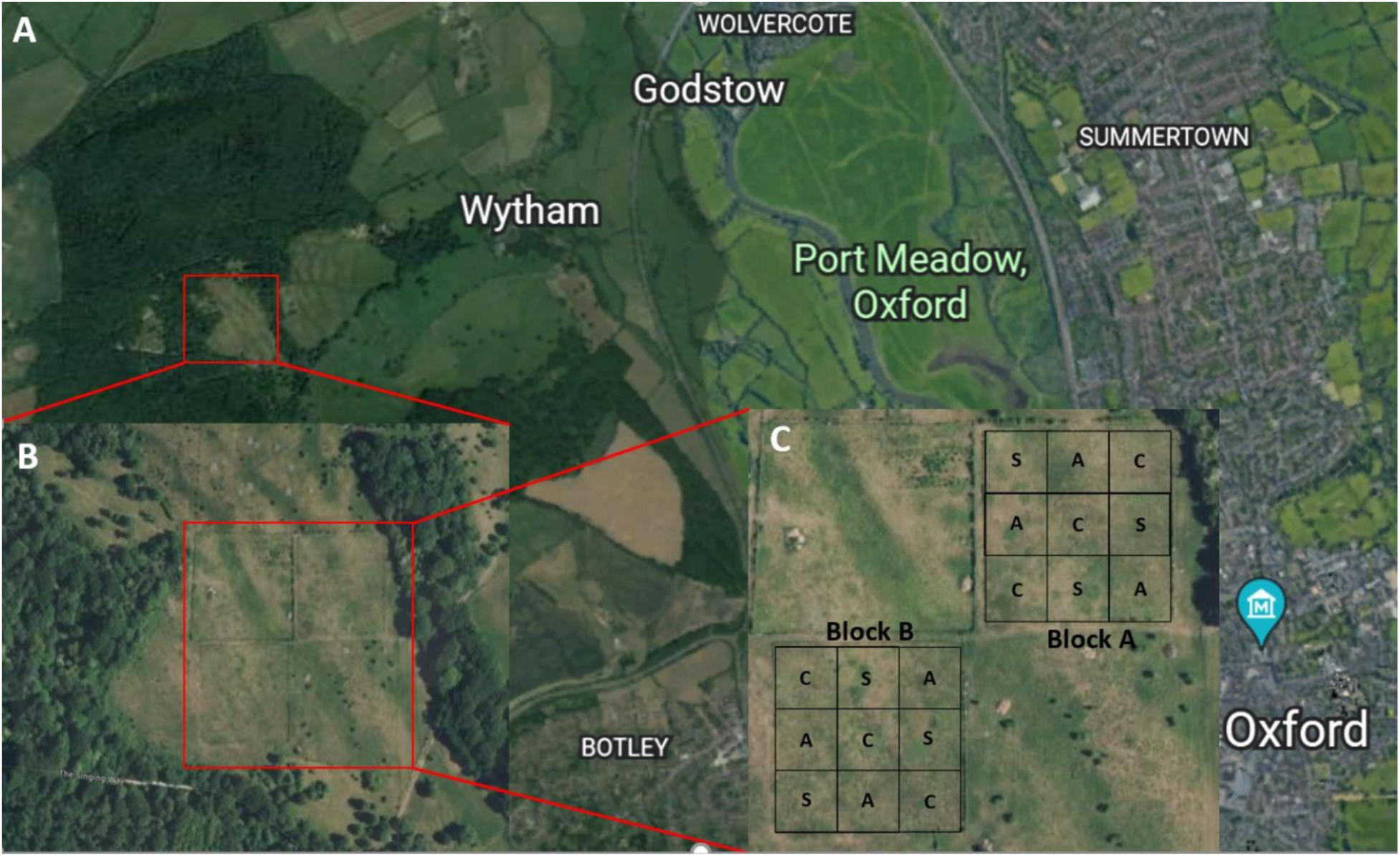
Study site: A) Map of Wytham Woods relative to central Oxford (5 km northwest of Oxford); B) Upper Seeds grassland within Wytham Woods. C) Gibson Grazing and Successional Experiment within Upper Seeds. The experiment has a replicated Latin grid square design with two blocks (A and B) and three treatments within each block (S = Spring-grazed, A = Autumn-grazed, C = Ungrazed control). Images retrieved from Google Satellite.

### Experimental design

The Gibson Experiment comprises three treatments imposed onto 18 30 x 30 m plots: two sheep grazed treatments (spring-grazed and autumn-grazed) and one ungrazed control. The grazing regime consisted of 3 sheep per paddock in each of the grazed plots. Because of the focus of the Gibson Experiment on successional dynamics the ungrazed plot was allowed to undergo secondary succession to a point at which young shrubs (mainly Hawthorn, *Crategus monogyna* Jacq.) approached a size at which they would be problematic to remove, at which point secondary succession was re-set by removal of shrubs and mowing. Hawthorn and other shrubs were removed to reset secondary succession in 2004, 2013 and 2017. The experiment has an unusual replicated blocked Latin square design with two blocks of nine plots and three replications of each treatment within each block (Figure 1C) (Gibson *et al*., 1992). This design was intended to encompass known gradients in soil quality across the field, with Latin squares orientated to maximise variation encompassed within control sites and thus increase confidence in any contrast detected with a grazing treatment. Thus, the total sample size is 18: 3 treatments (2 Grazed, 1 Ungrazed) x 3 replications of each treatment within each of two blocks (3 x 3 x 2 = 18 plots).

### Soil sampling

To compare grazed and ungrazed plots, we collected 18 composite samples (one from each of the 18 Gibson plots) from the top 5 cm of soil. Each composite sample consisted of 5 sub-samples (each 100 cm^3^) collected in a W-formation (Figure A1) using a soil core approximately 5 cm in diameter. Composite samples were thoroughly mixed and dried at 40°C to constant weight.

### Determination of soil physical and chemical properties

Soil pH was measured using a 1:1 soil-water ratio. Bulk density (BD) was calculated for each composite sample (500 cm^3^) as:

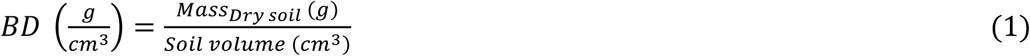

where the dry soil mass refers to the mass of the sample after it has been dried at 40°C. To correct for inorganic C-content (i.e., calcium-carbonate), we used the standard loss on ignition method, first heating the samples at 550°C to measure organic carbon and then at 950°C to obtain the mass of CaCO_3_. We calculated the inorganic carbon stocks in the top 5 cm of soil as follows (Equation 2), where 0.12 corresponds to the carbon mass fraction in CaCO_3_ (Soil Survey Staff, 2011):

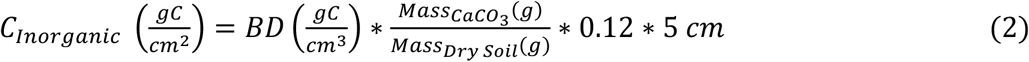

To estimate the total carbon and nitrogen content (%) of the samples, we performed CN elemental analysis using standard laboratory protocols (Soil Survey Staff, 2011). To divide the organic matter into mineral associated organic matter and particulate organic matter, we performed size-fractionation of organic matter. This method separates the soil into two size-fractions: >53 μm, which is dominated by particulate organic matter, and <53 μm, which is dominated by mineral-associated organic matter. To estimate the carbon and nitrogen content of each of the fractions in each of the samples, we performed CN elemental analysis as described above after size-fractionation. We calculated the carbon and nitrogen stocks in each fraction as follows (Equation 3), where carbon can be interchanged with nitrogen and mineral-associated organic matter with particulate organic matter.

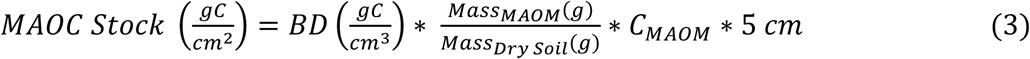

We calculated the C:N ratio in each fraction as follows (Equation 4), where mineral-associated organic matter can be interchanged with particulate organic matter.

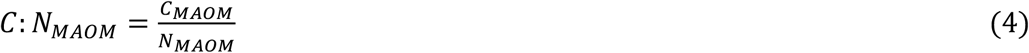

### Statistical analysis

Our analysis used linear mixed-effects models implemented with version 3.3.3 of the nlme R package in R (Pinheiro *et al.,* 2024) following a model building approach (Pinheiro & Bates, 2000). For this research question, the latin square design is overly complex relative to the limited sample size leading to singularities when trying to estimate the coefficients in complex models, necessitating simplification to achieve a model that would converge and produce credible estimates of the coefficients of interest. The simplified mixed-effects model included a fixed effect for the treatment of interest (grazing) and a random effect for blocks (estimates of treatment effects were very similar with alternative simpler models - see Appendix). A generic formula for the mixed effects model using the R language implementation of the Wilkinson and Rogers (1973) syntax is:

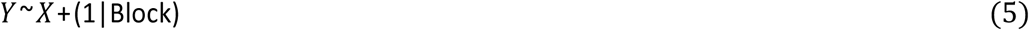

where, “Y” is a continuous response variable (one of the 12 listed in Table A2) and “X” is a fixed factor with either three levels (spring-grazed, autumn-grazed, ungrazed) or a simplified two-level factor comparing grazed and ungrazed plots. “Block” is a random factor with two levels for blocks (Gelman & Hill, 2007).

## Results

Of the 24 formal treatment comparisons only one was conventionally statistically significant (see below). The results should be interpreted considering the likelihood of one significant result for every twenty tests at the *P* < 0.05 level.

### Effects of grazing on soil properties

To address our main hypothesis on the effects of grazing we first performed comparisons of the soil properties in the grazed and ungrazed plots (i.e., pooling the spring and autumn grazed plots). In general, estimates of soil properties in the grazed and ungrazed treatments were similar with large overlap of confidence intervals (Table 1; Figure 4). The mixed-effects model analysis revealed that sheep grazing did not affect the mineral-associated organic carbon stock in the top 5 cm of soil (Figure 2). In addition, sheep grazing had no significant effect on the particulate organic matter carbon stocks, total organic carbon stocks, inorganic carbon stocks, or total carbon stocks, which all had relatively similar mean values and overlapping confidence intervals (Table 1; Figure 4; Table A3).

**Figure 2.**
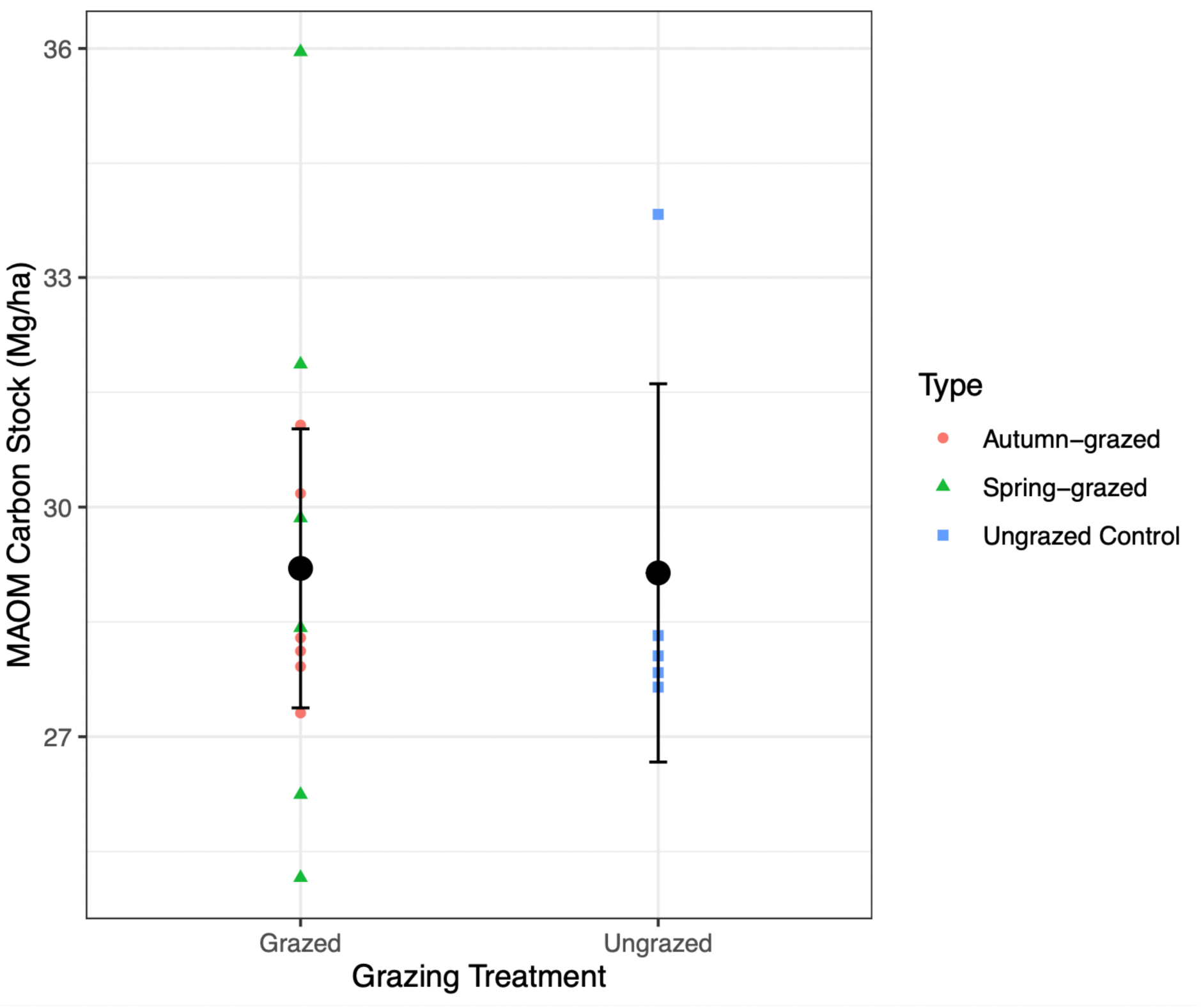
Carbon stocks in the MAOM fraction in grazed and ungrazed treatments: Small coloured points refer to individual plots. Larger black dots represent the treatment means and bars indicate the 95% confidence intervals from the linear mixed-effects model analysis likelihood profiles (*n_total_* = 18; *n_ungrazed_* = 6, *n_grazed_* = 12).

**Table 1.** Effects of grazing on selected soil properties in the top 5 cm. Treatment means with 95% CI bounds for ungrazed and grazed plots (combined and separately for spring and autumn grazed).

| Response | Ungrazed (*n* = 6) | Grazed (*n* = 12) | Spring-grazed (*n* = 6) | Autumn-grazed (*n* = 6) |
| --- | --- | --- | --- | --- |
| MAOM C stocks (Mg C ha<sup>-1</sup>) | 29.14 (26.67, 31.61) | 29.2 (27.38, 31.02) | 29.58 (25.44, 33.73) | 28.8 (27.27, 30.36) |
| POM C Stocks (Mg C ha<sup>-1</sup>) | 7.66 (6.19, 9.13) | 7.32 (6.22, 8.42) | 7.9 (5.59, 10.21) | 6.7 (5.73, 7.74) |
| SOC (Mg C ha<sup>-1</sup>) | 36.8 (33.69, 39.91) | 36.52 (33.82, 39.22) | 37.49 (31.27, 43.70) | 35.55 (33.92, 37.19) |
| Inorganic C (Mg C ha<sup>-1</sup>) | 4.77 (3.27, 6.27) | 5.16 (4.12, 6.20) | 5.65 (3.39, 7.90) | 4.68 (3.76, 5.59) |
| MAOM N Stocks (kg N m<sup>-2</sup>) | 0.25 (0.23, 0.28) | 0.24 (0.23, 0.25) | 0.24 (0.21, 0.26) | 0.24 (0.22, 0.27) |
| POM N Stocks (kg N m<sup>-2</sup>) | 0.016 (0.015, 0.018) | 0.015 (0.013, 0.017) | 0.015 (0.013, 0.017) | 0.014 (0.011, 0.018) |
| Total Organic N (kg N m<sup>-2</sup>) | 0.269 (0.245, 0.293) | 0.257 (0.243, 0.270) | 0.254 (0.229, 0.279) | 0.259 (0.239, 0.279) |
| MAOM C:N Ratio | 11.53 (11.06, 12.01) | 12.09 (11.67, 12.50) | 12.37 (11.57, 13.17) | 11.80 (11.37, 12.24) |
| POM C:N Ratio | 47.81 (38.32, 57.30) | 49.78 (42.80, 56.76) | 51.62 (38.32, 64.91) | 47.94 (37.61, 58.28) |
| SOC C:N Ratio | 14.40 (13.21, 15.59) | 15.52 (14.04, 16.70) | 16.35 (13.14, 19.57) | 14.68 (13.69, 15.68) |
| pH | 7.57 (7.46, 7.67) | 7.65 (7.58, 7.71) | 7.70 (7.62, 7.79) | 7.60 (7.49, 7.70) |
| Bulk density (g cm<sup>-3</sup>) | 0.78(0.71, 0.86) | 0.78 (0.73, 0.83) | 0.81 (0.72, 0.89) | 0.75 (0.67, 0.84) |

Across all treatments, the carbon stored in the mineral-associated organic matter fraction made up most of the total soil carbon pool (range: 53.4-89.1%). The particulate organic matter carbon stocks made up a smaller pool (range: 12.3-26.4%). The smallest but still sizeable pool was inorganic carbon (range: 7.9-19.8%) (Table 1; Figure 4). The C:N ratio of the particulate and mineral-associated organic matter fractions reflects how much nitrogen is required to sequester a unit of carbon within that fraction. Across all treatments, the C:N ratio was higher in the particulate fraction than in the mineral-associated organic matter fraction, as expected (Table 1). Furthermore, differences in C:N ratio within a fraction of organic matter reflect differences in organic matter formation and the efficiency of carbon sequestration within that fraction.

Sheep grazing led to a greater C:N ratio in the mineral-associated organic matter fraction compared to the ungrazed treatment (Table 1; Figure 4). Sheep grazing did not influence the C:N ratio in the particulate organic matter fraction or the total C:N ratio (Table 1; Figure 4). The sheep grazed and ungrazed treatments had similar total nitrogen stocks (Table 1; Figure 4; Table A3). Sheep grazing did not affect soil pH, with grazed and ungrazed plots having relatively similar mean values and overlapping confidence intervals (Table 1; Figure 4). Finally, the bulk densities in sheep grazed and ungrazed treatments were nearly identical, suggesting that sheep grazing did not affect bulk density (Table 1; Figure 4; Table A3).

### Effects of grazing type (timing) on soil properties

After performing an initial contrast of grazed versus ungrazed plots we proceeded to separate the spring and autumn grazed plots to compare all three treatments. Once again, soil properties were generally similar across treatments with overlap of confidence intervals. Grazing timing had a conventionally significant effect on only one soil property and a marginal effect on a second. First, grazing timing had a marginal positive effect on pH (Table A4). Spring grazing led to a higher (more alkaline) pH than the ungrazed control, but autumn grazing did not alter the pH relative to the ungrazed control. However, the absolute difference is small (although the logarithmic scale of the pH unit must be kept in mind): the pH in the spring-grazed plots was 0.13 pH units higher than the ungrazed control (Table 1). Second, grazing timing had a significant positive effect on the mineral-associated organic matter C:N ratio (Table 1, Table A4). Spring grazing produced the highest C:N ratio in the mineral-associated organic matter fraction (Table 1), significantly greater than the ungrazed control. In contrast, autumn grazing did not affect the C:N ratio in the mineral-associated organic matter fraction relative to the ungrazed control (Figure 3, Table A4).

**Figure 3.**
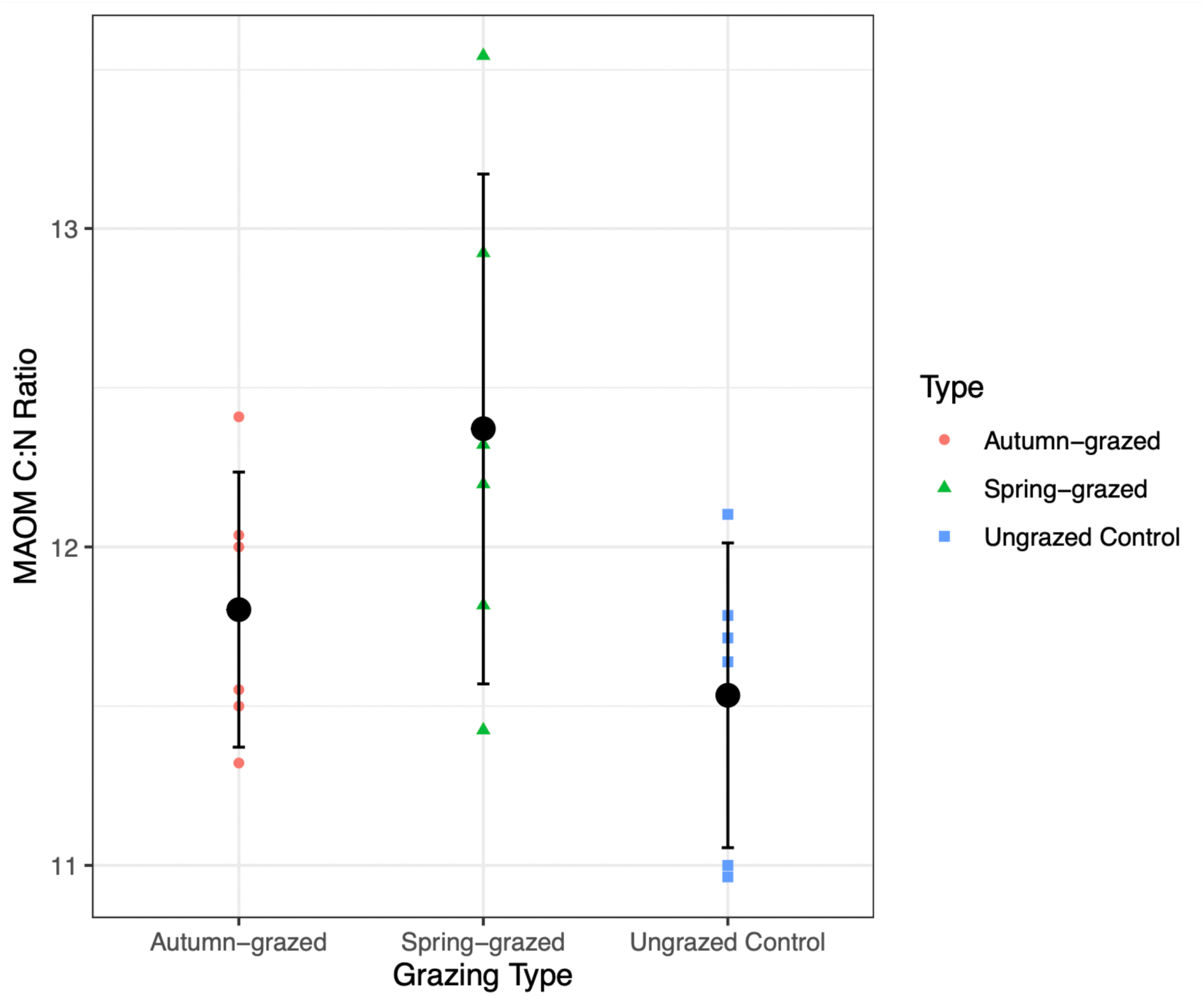
Comparison of C:N ratio in the MAOM of ungrazed, autumn-grazed, and spring-grazed plots: Small coloured points refer to the MAOC stocks found in the individual plots. Larger black dots represent the grazing type (timing) means and bars indicate the 95% confidence intervals from the linear mixed-effects model analysis likelihood profiles (n_total_ = 18; *n_treatment_* = 6).

## Discussion

We used a long-term (35 years) sheep grazing experiment to test the hypothesis that grazing increases soil carbon persistence by increasing the carbon stored in the more stable mineral-associated organic matter fraction. In this case, sheep grazing did not enhance the mineral-associated stocks (Figure 2). Furthermore, sheep grazing did not affect total carbon stocks (Table 1, Figure 4; Table S3). The only conventionally statistically significant result (p < 0.05) from the 24 formal tests performed was of higher mineral-associated organic matter C:N ratio in spring-grazed plots (Figure 3, Table A4). Since one false positive result is expected at this significance level for every 20 tests performed this result could be a false positive and needs confirming by independent study.

**Figure 4.**
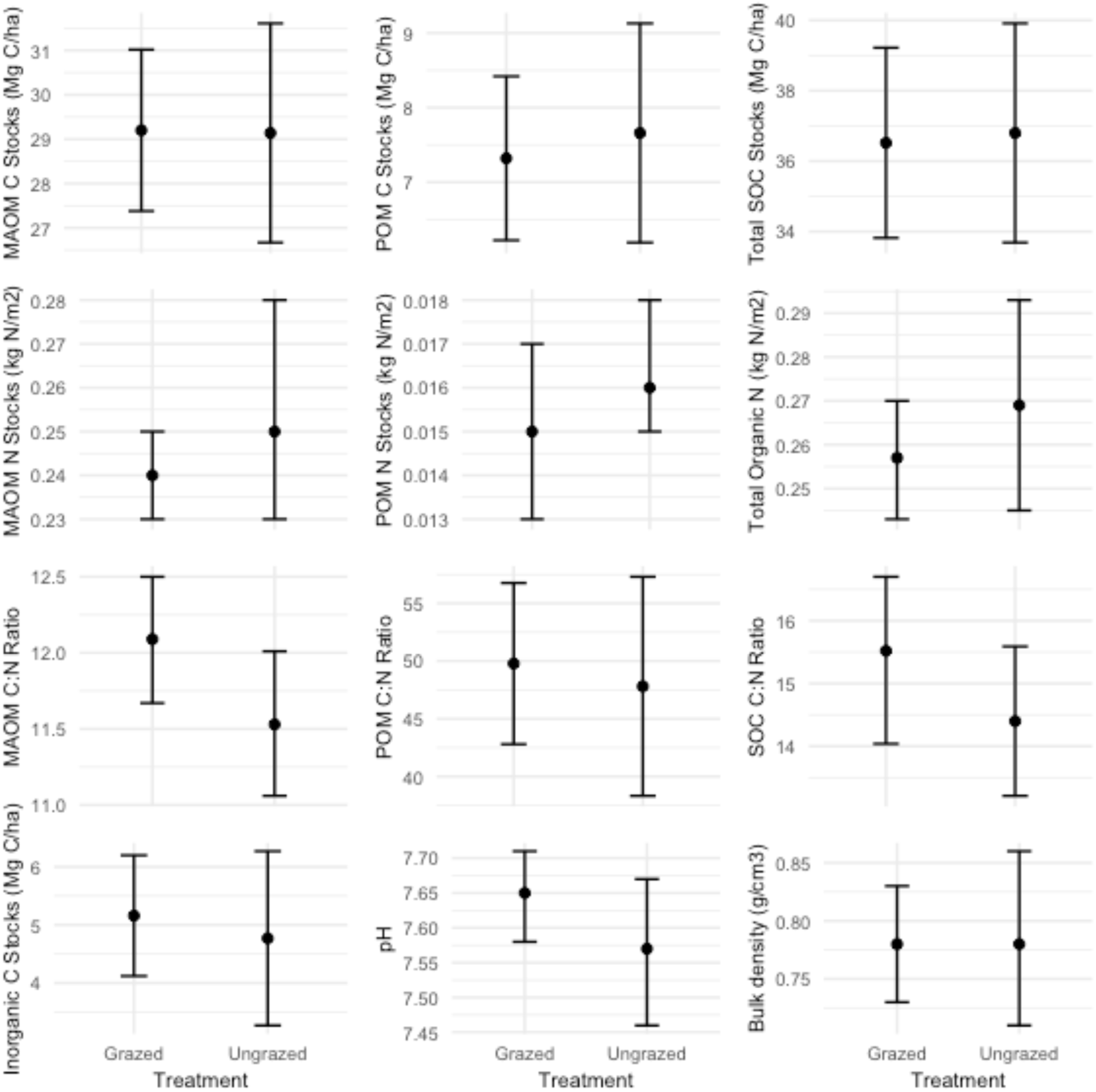
Effects of grazing on selected soil properties. Black dots represent the treatment means and bars indicate 95% confidence intervals from the linear mixed-effects model analysis likelihood profiles.

### Sheep grazing did not affect soil carbon sequestration or persistence

In this study, sheep grazing did not increase mineral-associated carbon sequestration (Figure 2). This suggests that sheep grazing does not lead to a more persistent soil carbon pool in this system, contrary to the hypothesis proposed by Kristensen *et al*. (2022). This could be explained by the neutral effect of sheep grazing on soil nitrogen stocks and pH, the saturation of mineral surfaces, and/or shallow sampling depth. Kristensen *et al*. (2022) suggested that grazing by large herbivores should enhance mineral-associated carbon sequestration in part by increasing soil fertility (particularly nitrogen), a key determinant of soil carbon sequestration. Increased fertility is expected to increase plant productivity and by extension increase the quantity of organic matter inputs to the soil. As shown above, this grazing regime did not increase soil nitrogen stocks, which could explain why it did not lead to greater overall carbon storage.

The persistence of soil carbon is also highly nitrogen-dependent, with mineral-associated organic matter generally having a lower C:N ratio than particulate organic matter (Lugato *et al*., 2021). This general trend is reflected in this calcareous grassland as well, with the particulate organic matter C:N ratio higher than the mineral-associated organic matter C:N ratio across all treatments (Table 1). Grazing is hypothesized to increase soil nitrogen stocks, but our results suggest that this grazing regime did not increase the amount of nitrogen stored in the soil organic matter (Table 1; Figure 4; Table S3). This could be due to the “harvest export” (nutrient stripping) from the system, where the sheep are moved to different fields (and ultimately from the system when they are used for products such as wool and meat) although they may also import nutrients when moved into an area. The fact that in this case sheep grazing produced no change in soil organic nitrogen stocks could explain why there was no increase in mineral-associated carbon stocks.

It is important to note that all plots experience encroachment by scrub (Hawthorn etc.) over time (spring and autumn grazing only were insufficient to prevent scrub build up) necessitating three rounds of removal (see Methods) with potential export of nutrients in the vegetation. However, scrub build up was more intense in the ungrazed than grazed plots and it is unlikely that the unequal removal of scrub across the treatments would act in a way to equalize soil characteristics.

Sheep grazing also did not affect bulk density. Trampling by grazers is often expected to increase bulk density by compacting the soil. However, in this case, bulk density was uniformly low (Table 1; Figure 4), perhaps because the grazing treatments in this study involved short duration, low intensity (3 sheep per paddock) grazing that did not compact the soil beyond its natural resilience, as sheep tend to compact soil less than larger grazers like cattle (Cournane *et al*., 2011; Gibson (2011)).

### Saturation of mineral surfaces

In this study, mineral-associated organic matter made up a high proportion of soil organic carbon and total soil carbon (53.4-89.1% across all treatments) (Table 1). This suggests that the soils at Upper Seeds may have approached their finite mineral-associated carbon storage capacity. Mineral-associated carbon sequestration is saturating, whereas particulate carbon sequestration is not. The amount of carbon that can be stored in the mineral-associated organic matter is finite and largely determined by the availability of adsorption sites at mineral surfaces, hence effectively a result of the clay content and type (Georgiou et al. 2022). This means that carbon inputs to the soil can only be incorporated into mineral-associated organic matter when there is a “saturation deficit” (Castellano *et al*., 2015). Once the mineral surfaces become saturated with carbon, mineral-associated carbon stocks can no longer increase, meaning that the organo-mineral exchange is at dynamic equilibrium (Cotrufo *et al*., 2019). Thus, it is possible that the lack of effect of sheep grazing on mineral-associated carbon stocks shown here reveals that this soil has become carbon-saturated and that there are no available mineral surfaces to shield the organic matter from decomposition. Recently, it was shown that mineral surfaces in natural grasslands are closer to saturation than in managed grasslands (Georgiou et al. 2022). Hence, our results may suggest that the management regime at Upper Seeds was too extensive to significantly shift the soil dynamics away from their natural state. Further analyses such as assessing the grain size distribution and mineralogy in the soil could serve as proxies for the sorption capacity of the soils, which could be used to determine if the soil has reached its finite sorption capacity (Abramoff *et al*., 2021).

### Sheep grazing did not reduce soil pH

Changes in soil pH as a response to grazing has been shown in previous studies (e.g., Hiernaux *et al.,* 1999). The narrow pH range in our soils showed that grazing had little impact on soil acidity at Upper Seeds, and hence no pH-induced influence on mineral-associated organic matter sorption rate and capacity (Abramoff et al. 2021). In a shallow calcareous grassland with typically <20 cm from the surface to the calcareous bedrock, the adsorption complex is expected to be close to saturation, particularly with base cations released from limestone weathering. This gives a high acidity buffer capacity, which is in line with our pH values well above neutrality in the very top of the soil subjected to the highest degree of acidification.

### Herbivore diversity

The lack of herbivore diversity could explain the observed neutral effect of sheep grazing on soil carbon stocks. Overall, herbivore assemblage determines what plants are preferentially consumed, which influences the plant community composition and by extension the input of organic matter to the soil (Chang *et al*., 2018). Several studies (e.g., Sitters *et al*., 2020; Li *et al*., 2021) suggest that grazing by a single herbivore species may even decrease soil carbon stocks. This is likely due to specific dietary preferences, foraging modes and trampling effects that push the plant community towards dominance by a few well-adapted species which may have reduced belowground carbon inputs. This is particularly the case for sheep grazing. Sheep, which are smaller and more selective than some large herbivores like cattle, tend to feed on highly nutritious forb species, which reduces their abundance in grassland communities and their production of organic matter inputs. In contrast, larger herbivores such as cattle tend to be less selective and can create a more even plant community (Chang *et al*., 2018). This suggests that the presence of multiple functional groups of herbivores (e.g., cattle and sheep) may counteract the effects of sheep grazing alone. Combining multiple different herbivore species leads to a greater diversity of foraging modes, dietary selections, and trampling effects, which in some cases can maintain more diverse plant communities with more consistent carbon inputs (Li *et al*., 2021). Here, the presence of only a single herbivore species (as is typical in most livestock systems) may be limiting the realized soil carbon storage. However, it is also important to emphasize that agricultural grazing systems tend to use single species and so the case examined here is typical in this regard.

### Timing of grazing

The mineral-associated organic matter C:N ratio was significantly higher in the sheep grazed treatment than in the ungrazed control. However, this was not consistent across the two grazing times, with only spring-grazing leading to a higher mineral-associated C:N ratio than the ungrazed control (Figure 3, Table A4). The C:N ratio can be considered a fingerprint of what types of carbon inputs end up forming mineral-associated organic matter. As the C:N ratio decreases during decomposition, the higher C:N ratio in the spring-grazed plots suggests that a higher proportion of the organic matter in the mineral-associated fraction in the spring-grazed treatment is formed directly from dissolved organic carbon from plants (e.g., root exudates) rather than microbial necromass or shoot tissue. Dissolved organic carbon from rhizodeposits have a higher average C:N ratio than other carbon inputs such as shoot tissues (Ostrowska & Porebska, 2015). Thus, the higher C:N ratio in the spring-grazed treatment could suggest that spring-grazing leads to greater rhizodeposition, probably due to compensatory growth (Villarino et al. 2021; Hamilton et al 2008). Compensatory growth is a strategy evolved to cope with disturbances causing a loss of biomass. When plants are consumed by grazers, they compensate by scavenging for nutrients and extending their root systems to support regrowth. This often involves increasing root exudation in exchange for nutrients from soil microorganisms but is only relevant during periods of active plant growth. Hence, in the early growing season, when both growth rates and forage quality are highest, this mechanism is most pronounced. This could explain why the mineral-associated organic matter C:N ratio is higher in spring-grazed but not in autumn-grazed plots.

### Caveats and limitations

Our study site at the top of Wytham hill on a bedrock of coral rag limestone has shallow soils, especially following the ploughing for agricultural use between the 1950s and 80s. Generally, the proportion of mineral-associated organic matter relative to particulate organic matter increases with depth such that the results observed here may not extend to deeper soil.

The cessation of the continuous management of the Gibson Grazing and Successional Experiment in 2020 and subsequent removal of the fences between the plots meant that sheep were free to graze the whole experiment for a short time before the soil sampling occurred. It seems unlikely that such a short time of free grazing would have eliminated any treatment effects that had built up over the 35 years of the Grazing Experiment.

## Author Contributions

DE conceptualized and designed the study with the help of AH and JK; DE collected the field data with help from AH and JK; DE conducted the laboratory and statistical analysis with the input from RB, DA, and ME; DE wrote the paper with input from all co-authors particularly AH and JK. CH helped managed the Grazing Experiment from 2013 to its cessation. All authors commented on and approved the manuscript.

## Acknowledgments

Nigel Fisher and team manage the Wytham Woods research estate, including the Upper Seeds grassland. JAK was supported by the Carlsberg Foundation (grants CF20_0238 & CF23_0641). The Gibson Grazing experiment was supported by the Ecological Continuity Trust, the Patsy Wood Trust and the British Ecological Society. AH was supported by the John Fell Fund.

## Data Availability Statement

The data that support the findings of this study are available from the corresponding author upon reasonable request.

## Appendix

### Statistical analysis

The blocked latin square design of the Gibson grazing experiment is too complex given the small sample size of 18 plots. A classical linear model (i.e. implemented with the R lm() function) that includes fixed factors for block and the rows and columns of the latin square design is over-parameterized such that some factor levels cannot be estimated. Mixed-effects models can be useful for the analysis of small data sets like this since treating factors as random effects requires fewer degrees of freedom because a single variance component (costing one degree of freedom) is estimated for each factor regardless of the number of levels. However, for this experimental design a mixed-effects ANOVA with random effects for block, row and column (fitted with the R lme4 package lmer() function) warns of singularities and contains estimates of zero for variance components for both block and column (see Supplementary R Markdown [pp. 10-14]). There was therefore no option but to fit simpler mixed-effects model: the small sample size was able to support a reduced model with a fixed effect for treatment and a random effect for block as reported in the main text (fitted using the lme() function from the nlme package for R). Importantly, the coefficients of interest – the treatment means (or differences in means) and their confidence intervals – were virtually unchanged by the different model formulations and so our results are insensitive to these fine details of model formulation.

**Figure A1.**
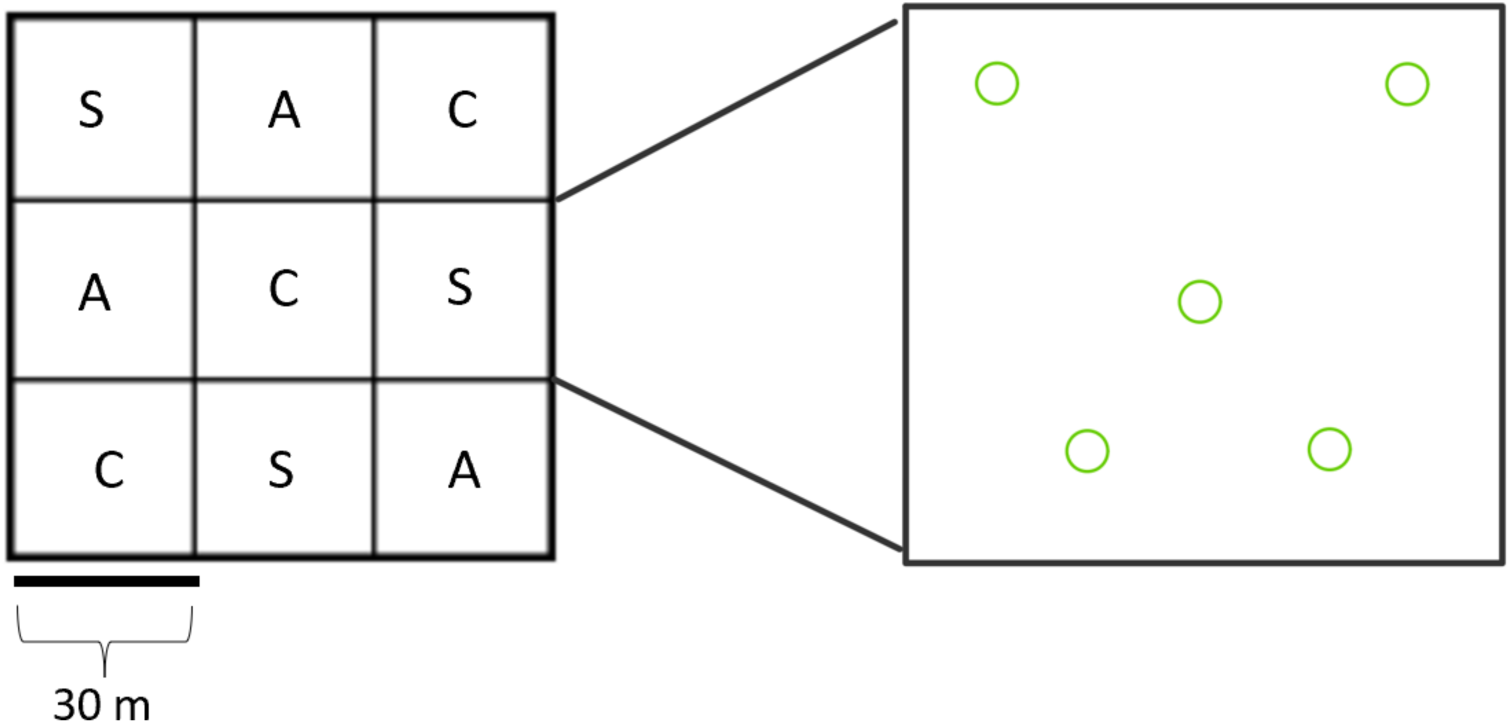
**One block of plots showing latin square layout** (Left: S = Spring-grazed, A = Autumn-grazed, C = Ungrazed control**) and the W-formation of sampling points within each plot** (Green circles refer to each of the 5 sub-samples taken within each plot. The 5 sub-samples within each plot together form one composite sample).

**Table A1.** Table of Abbreviations.

|  |  |
| --- | --- |
| CO<sub>2</sub> | Carbon dioxide |
| POM (POC) | Particulate organic matter (Particulate organic carbon) |
| MAOM (MAOC) | Mineral-associated organic matter (mineral-associated organic carbon) |
| SOC | Soil organic carbon |
| SOM | Soil organic matter |
| TOC | Total organic carbon |
| TOM | Total organic matter |
| N | Nitrogen |
| BD | Bulk density |
| C | Ungrazed control plots |
| S | Spring-grazed treatment |
| A | Autumn-grazed treatment |

**Table A2.** Response variables included in the statistical analysis.

| Response variable | Description |
| --- | --- |
| MAOM carbon stocks | Amount of carbon (Mg C/ha) stored in the MAOM fraction in the top 5 cm of soil |
| POM carbon stocks | Amount of carbon (Mg C/ha) stored in the POM fraction in the top 5 cm of soil |
| Total organic carbon stocks | Amount of carbon (Mg C/ha) stored in both the MAOM and POM fractions in the top 5 cm of soil |
| Inorganic carbon stocks | Amount of carbon (Mg C/ha) stored in the inorganic fraction in the top 5 cm of soil |
| Total carbon stocks | Amount of carbon (Mg C/ha) stored in both organic and inorganic carbon stocks in the top 5 cm of soil |
| MAOM nitrogen stocks | Amount of nitrogen (kg N/m<sup>2</sup>) stored in the MAOM fraction in the top 5 cm of soil |
| POM nitrogen stocks | Amount of nitrogen (kg N/m<sup>2</sup>) stored in the POM fraction in the top 5 cm of soil |
| Total nitrogen stocks | Amount of nitrogen (kg N/m<sup>2</sup>) stored in both the MAOM and POM fractions in the top 5 cm of soil. Based on the assumption that most soil nitrogen is stored in an organic form (Post *et al*, 1985). |
| MAOM C/N Ratio | Ratio of carbon to nitrogen in the MAOM fraction in the top 5 cm of soil |
| POM C/N Ratio | Ratio of carbon to nitrogen in the POM fraction in the top 5 cm of soil |
| Total C/N Ratio | Ratio of carbon to nitrogen in the top 5 cm of bulk soil |
| pH | Measure of acidity/alkalinity of the samples |
| Bulk Density | The mass of soil divided by the volume of the soil |

**Table A3.**
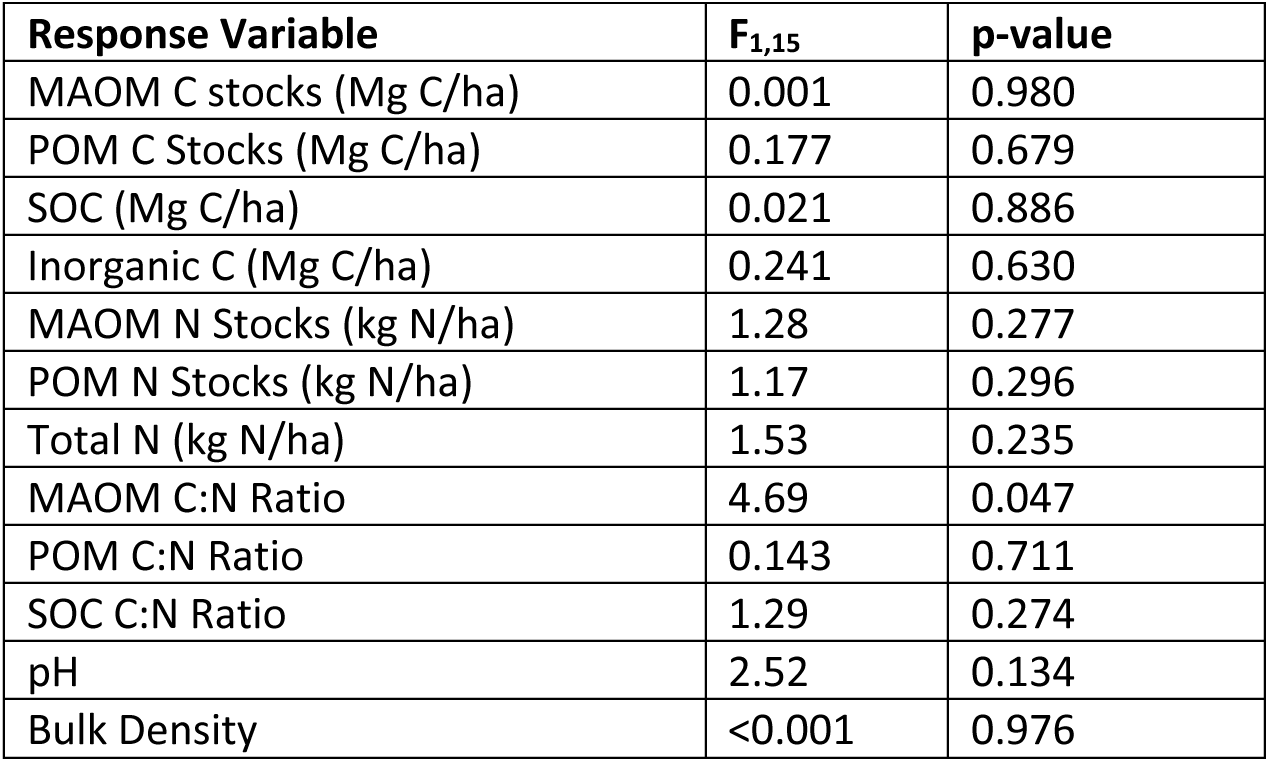
Grazing treatment (grazed vs ungrazed) ANOVA table.

**Table A4.** Grazing type (spring, autumn, ungrazed) ANOVA table.

| **Response Variable** | **F<sub>2,14</sub>** | **p-value** |
| --- | --- | --- |
| MAOM C stocks (Mg C/ha) | 0.092 | 0.912 |
| POM C Stocks (Mg C/ha) | 0.885 | 0.435 |
| SOC (Mg C/ha) | 0.330 | 0.724 |
| Inorganic C (Mg C/ha) | 0.693 | 0.517 |
| MAOM N Stocks (kg N/ha) | 0.728 | 0.500 |
| POM N Stocks (kg N/ha) | 0.773 | 0.480 |
| Total N (kg N/ha) | 0.812 | 0.464 |
| MAOM C:N Ratio | 4.93 | 0.024 |
| POM C:N Ratio | 0.247 | 0.784 |
| SOC C:N Ratio | 1.72 | 0.215 |
| pH | 3.56 | 0.056 |
| BD | 1.30 | 0.305 |

